# stMCP: Spatial Transcriptomics with a Model Context Protocol Server

**DOI:** 10.64898/2026.03.11.710153

**Authors:** Jordan J. Smith, Xi Wang, Matthew McPheeters, Made Airanthi Widjaja-Adhi, Sejiro Littleton, Daniel R. Saban, Marcin Golczak, Michael W. Jenkins

**Affiliations:** Department of Biomedical Engineering, Case Western Reserve University, Cleveland, OH, USA; Department of Pharmacology, Case Western Reserve University, Cleveland, OH, USA; Department of Ophthalmology, Duke University School of Medicine, Durham, NC, USA; Department of Integrative Immunobiology, Duke University School of Medicine, Durham, NC, USA; Department of Pediatrics, Case Western Reserve University, Cleveland, OH, USA; FES Ctr., Louis Stokes Cleveland VA Med. Ctr., Cleveland, OH

## Abstract

Spatial transcriptomics enables high-resolution mapping of gene expression in intact tissues but remains challenging due to complex computational workflows that limit accessibility and reproducibility. Here, we present a Model Context Protocol (MCP) framework enabling natural language-driven spatial transcriptomics analysis. By executing analytical tools locally, this architecture eliminates the need to upload massive datasets to large language models, bypassing high token costs and mitigating data privacy and training risks. The MCP orchestrator interprets intent, dynamically routes requests, maintains session state, and verifies input integrity to ensure reproducible execution. Benchmarking across biological discovery, orchestration accuracy, token usage, and execution time demonstrates robust performance. This architecture establishes a scalable template for AI-native research by standardizing the interface between models and local analytical engines. Rather than replacing bioinformaticians, this framework empowers biologists to independently and comprehensively explore their data, accelerating hypothesis testing, and unlocking broader biological discoveries.

## Main

Spatial transcriptomics technologies enable high resolution mapping of gene expression within intact tissues, providing insight into tissue organization, cellular interactions, and spatial patterns.^1–4^ Recent advances have substantially expanded the scale and resolution of spatial transcriptomics experiments, allowing simultaneous measurement of thousands of genes across many cells within a single tissue section.^5–10^ These approaches have been applied across diverse biological systems to study development, tissue homeostasis, and cancer, among others.^11–13^

Despite rapid technological progress, the analysis of spatial transcriptomics data remains challenging. Spatial datasets are inherently high dimensional, combining gene expression profiles with spatial coordinates and tissue morphology. Extracting meaningful biological insight typically requires iterative, exploratory analysis involving preprocessing, dimensionality reduction, clustering, spatial visualization, and spatial statistics. These workflows are further complicated by the need to integrate prior biological knowledge such as marker genes, anatomical context, and pathway knowledge, while remaining open to unexpected or rare biological signals.^14–16^

Current computational approaches to spatial transcriptomics analysis present significant bottlenecks. While manual coding workflows using toolkits such as Seurat, Scanpy, and Squidpy offer flexibility and methodological rigor, they require programming expertise that many biologists lack.^17–19^ Consequently, researchers must rely on bioinformaticians to execute and adjust scripts. This back-and-forth communication creates a slow, iterative loop that disrupts the natural flow of biological discovery, hindering the rapid, spontaneous hypothesis testing required to uncover novel insights. Conversely, automated pipelines streamline analysis but may constrain exploration and bias results toward dominant cell populations or predefined analytical paths, increasing the likelihood of overlooking certain aspects of the data. Because spatial transcriptomics analysis is rarely linear, requiring dynamic continuous interplay between visualization, statistical testing, and biological interpretation, these delayed or rigid workflows ultimately limit the depth of data exploration.

Large language models (LLMs) have recently emerged as powerful tools for translating natural language instructions into structured computational tasks, raising interest in their potential application to biological data analysis.^20^ In principle, LLMs could lower technical barriers by allowing researchers to specify analytical intent in natural language rather than code; however, there are significant concerns that include lack of transparency, prohibitive token costs, data privacy risks associated with uploading massive datasets, and the potential for hallucinated results.^21^ These limitations have motivated calls for constrained approaches in which language models assist with orchestration and interpretation rather than performing analytical computation directly.^22,23^

Recent efforts in spatial and single-cell transcriptomics have begun to reconcile natural language reasoning with domain-specific data analysis through agentic and foundation model frameworks. Autonomous AI agents such as *SpatialAgent* integrate large language models with dynamic tool execution to automate multistep spatial biology workflows spanning experimental design, multimodal data analysis, and hypothesis generation.^24^ Similarly, *STAgent* is a multimodal LLM-based AI agent tailored for spatial transcriptomics that can generate code, interpret spatial patterns, and synthesize scientific reports with minimal human input.^25^ Complementing these agentic workflows, domain-adapted foundation models like *scGPT* extend pretrained single-cell transformers to spatial transcriptomics through large-scale training on transcriptomic data, enabling improved cell type annotation, batch correction, and multi-omic integration.^26^ While these approaches demonstrate automated reasoning and native integration with analytical toolchains, they remain dependent on LLM planning and interpretation layers and, as a result, continue to inherit the nondeterminism, limited interpretability, and high cost of general language models.

Much as HTTP became the universal standard for data exchange on the web, the Model Context Protocol (MCP) is poised to become the foundational standard for agentic tool use. MCP is an open standard designed to bridge this gap through a secure client-server architecture that connects AI models to external tools and datasets.^27,28^ In an MCP-based workflow, the language model is restricted to interpreting user intent and issuing replies to the user, while all numerical computation is executed locally by validated analytical software. This separation reduces reliance on model-generated code and avoids uploading raw biological data to remote services. Moreover, because MCP tool modules reside on a local server with explicitly defined interfaces, the analytical pipeline is insulated from the “brittle API” problem, where upstream changes typically disrupt agent-based systems. Updates to analytical capabilities occur within the server environment itself, allowing version-controlled tool evolution without compromising workflow stability. By encapsulating analytical functions within deterministic server modules and comprehensively logging tool calls, parameters, and outputs, MCP enables flexible multistep exploration while preserving reproducibility, transparency, and cost-efficient operation.

Here, we present an MCP framework specifically tailored for spatial transcriptomics analysis and apply it to Xenium data from mouse corneal tissue to demonstrate its practical and biological utility. We show that structured, iterative querying through natural language enables the rapid identification of rare, low-abundance Schwann cell populations, as well as the resolution of specific immune cell subsets within the tissue microenvironment, that are not readily apparent in standard, fixed clustering outputs. To validate technical performance, we evaluated the system using a predefined prompt framework, rigorously benchmarking orchestration accuracy, language model token consumption, and local execution time. Collectively, our results establish this MCP framework as a robust, assistive computational tool that empowers biologists to comprehensively explore their data while preserving methodological rigor, data governance, and reproducibility.

## Results

We developed a Model Context Protocol (MCP) server that pairs locally executed analytical tools with the contextual reasoning and domain knowledge of large language models, enabling reproducible spatial transcriptomics analysis driven entirely by natural language. We evaluated the MCP server using a mouse cornea dataset generated via the Xenium spatial transcriptomics platform to demonstrate efficient token usage, reproducible analysis workflows, and seamless integration of preprocessing, visualization, and spatial statistics. We also show that using MCP can help find subtle information in the dataset that may be hard to detect when following normal protocols.

### System Overview and Design Principles

The spatial transcriptomics MCP framework is designed to enable natural language-driven analysis while ensuring reproducibility, transparency, and data privacy through local execution. The system architecture is organized into three primary, interacting components: an LLM-powered MCP host, a local MCP server (comprising an orchestrator and analytical tools), and a web application for showing results (Fig. 1).

**Figure 1:**
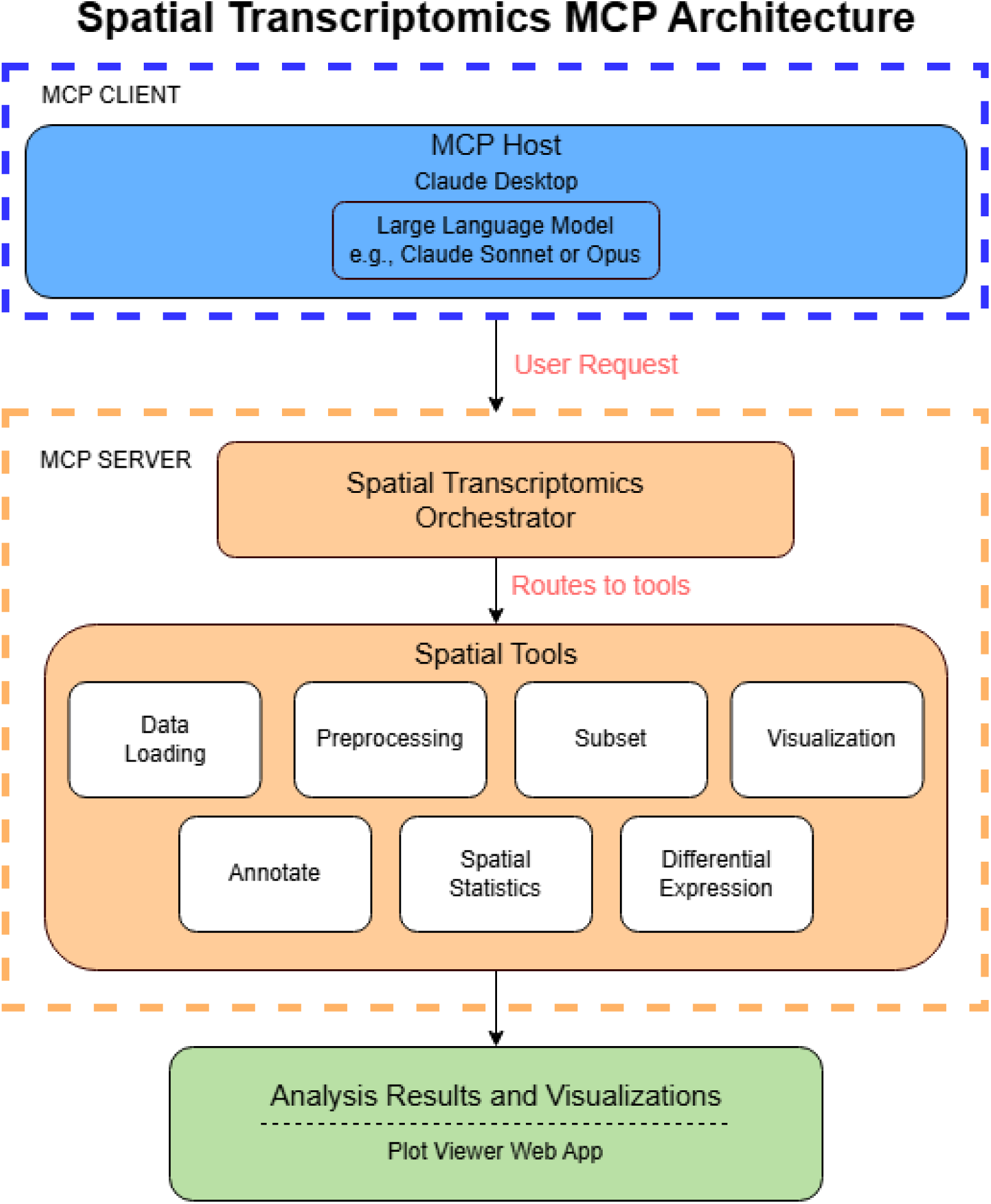
Overview of the MCP framework for spatial transcriptomics. User prompts are issued through the MCP client (LLM (e.g., Claude Opus) and sent to the MCP server, which consists of an orchestrator and modular spatial analysis tools. The orchestrator routes structured requests to the tools and manages session state. Outputs, including visualizations and data objects, are returned to the user and logged for reproducibility.

#### The MCP Host and LLM Interface

The MCP host serves as the user-facing interface, translating natural language queries into structured analytical intent. While the framework is fundamentally LLM-agnostic and can interface with any compatible model, our current implementation utilizes Claude Desktop as the host, leveraging advanced models (e.g., Sonnet and Opus) (Fig. 2). Crucially, the LLM is restricted strictly to interpreting user queries, generating structured tool requests, and summarizing results (Supplementary Fig. 1). This architectural separation ensures that the LLM performs no numerical computation, data manipulation, or independent code execution, thereby preventing hallucinations during data processing.

**Figure 2:**
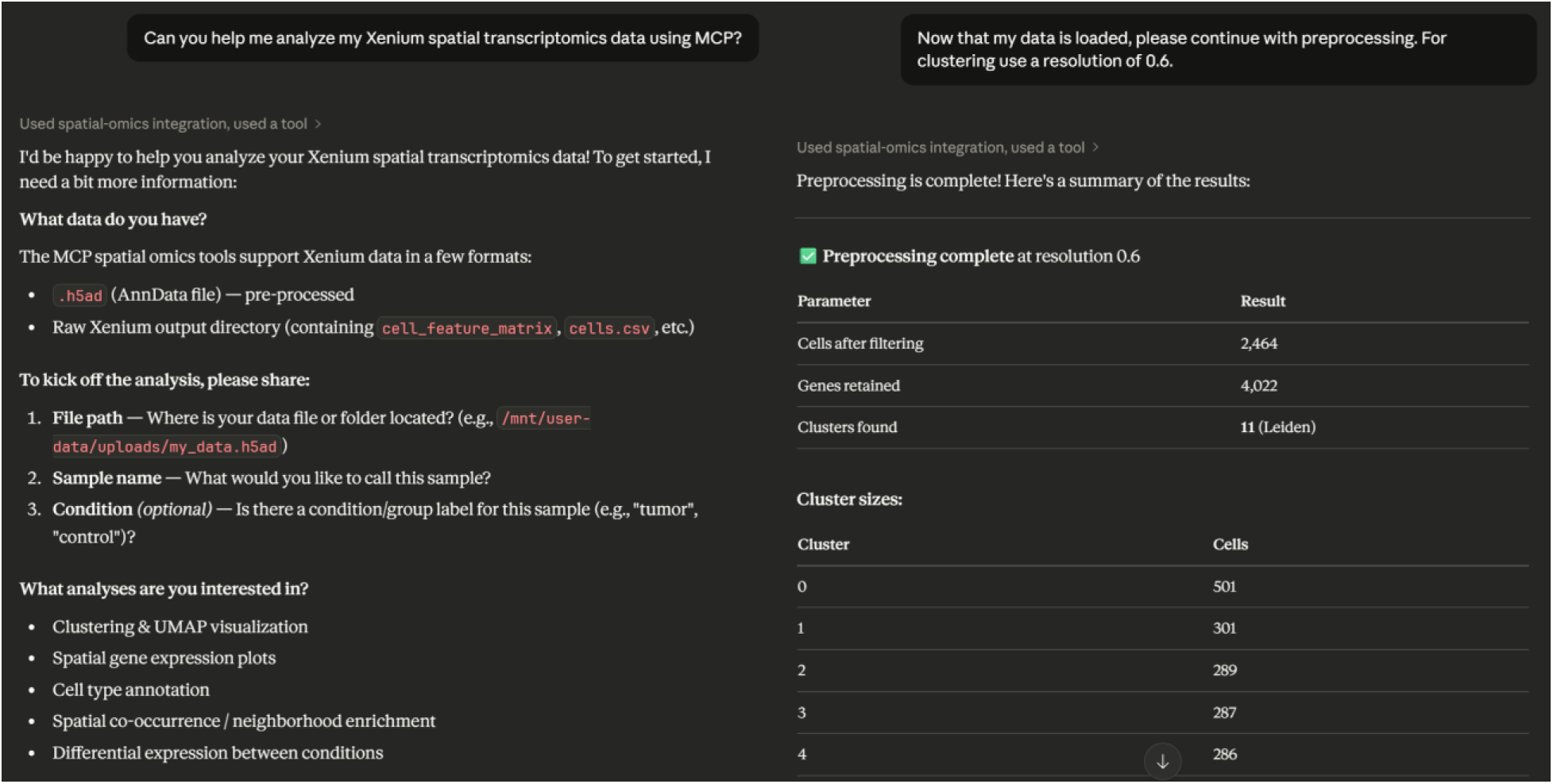
End-user interface of the stMCP. Screenshot of the Claude Desktop application connected to the MCP server, illustrating a natural language query and the corresponding execution.

#### The Local MCP Server - Orchestration, Session Management, and Logging

The core computational engine is the local MCP server, structured as a two-tiered system containing an orchestrator layer and a suite of local analysis tools. The orchestrator acts as a supervisory control layer; it receives structured requests from the host and mediates communication with the downstream tools.

To ensure continuity across multistep workflows, tool execution is coordinated through a shared session context managed by the orchestrator. This abstraction maintains references to the active dataset, output directory, tool state, and analysis history, preventing unintended state inconsistencies between tool calls. Furthermore, all MCP actions, including user prompts, validated tool invocations, parameter specifications, and generated outputs, are automatically recorded in a structured metadata file. This comprehensive logging framework captures both analytical intent and computational execution, creating a complete audit trail that allows workflows to be reproduced, inspected, and programmatically reconstructed (Supplementary Fig. 2).

#### Tool Module Implementation and Modular Design

Once validated by the orchestrator, analytical operations are executed locally as independent tool modules (Table 1). The MCP tool suite spans the core components of a spatial transcriptomics analysis pipeline, including data loading, data subsetting, preprocessing, clustering, visualization, annotation, spatial statistics, and differential analysis. The MCP can load in data from Xenium, Vizgen MERSCOPE, and CosMx Nanostring, as well as any Anndata object.^3,29–31^

**Table 1:**
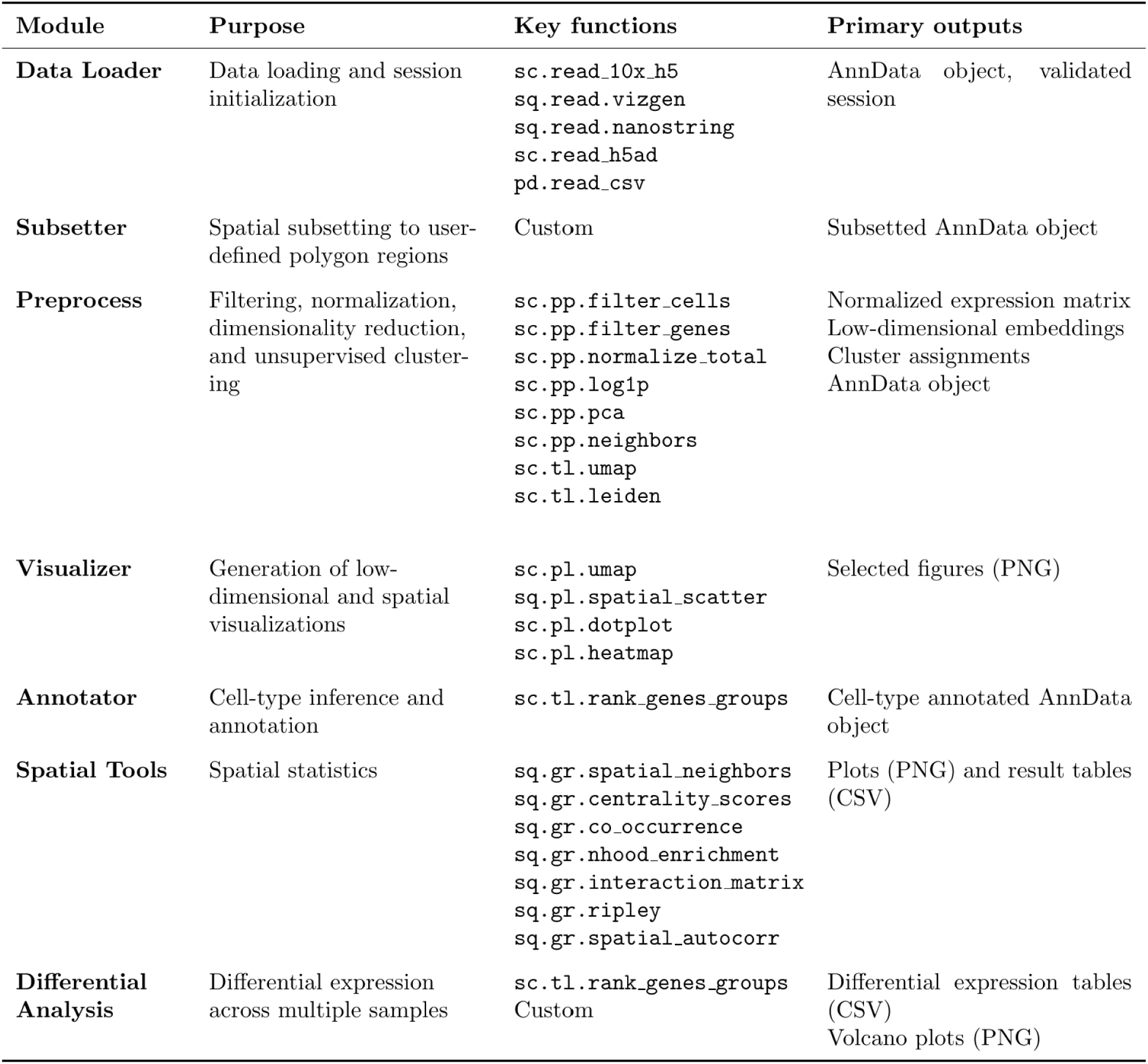
Spatial Transcriptomics MCP Tools Overview. This includes the tool name, the task, its basic Python functions, and the expected outputs for each tool. The key functions use sc for scanpy, sq for squidpy, and pd for pandas.^18,19,32^

These tools leverage established Python libraries, including Scanpy and Squidpy, to ensure methodological rigor and deterministic results directly comparable to manual workflows (Supplementary Fig. 3).^18,19^ All tools conform to a unified function interface with explicitly defined input and output parameters. Users can easily override default parameters via natural language to tailor the analysis to their specific experimental needs. Importantly, this modular architecture supports seamless extensibility; new analytical capabilities can be incorporated as additional modules without modifying the underlying orchestrator’s logic. Tool wrapping is simple with an orchestrator, tool module, and function layers (Supplementary Fig. 4).

#### Web-Based Results Viewer

To complete the exploratory loop, the framework integrates a dedicated web application for the immediate inspection of analytical outputs. Rather than requiring users to manually navigate the local file system, the viewer provides a centralized interface to browse and review all generated outputs. The viewer automatically detects new files produced within the user-defined output directory, tightly coupling tool execution with immediate feedback to support an intuitive, iterative discovery process (Fig 3).

**Figure 3:**
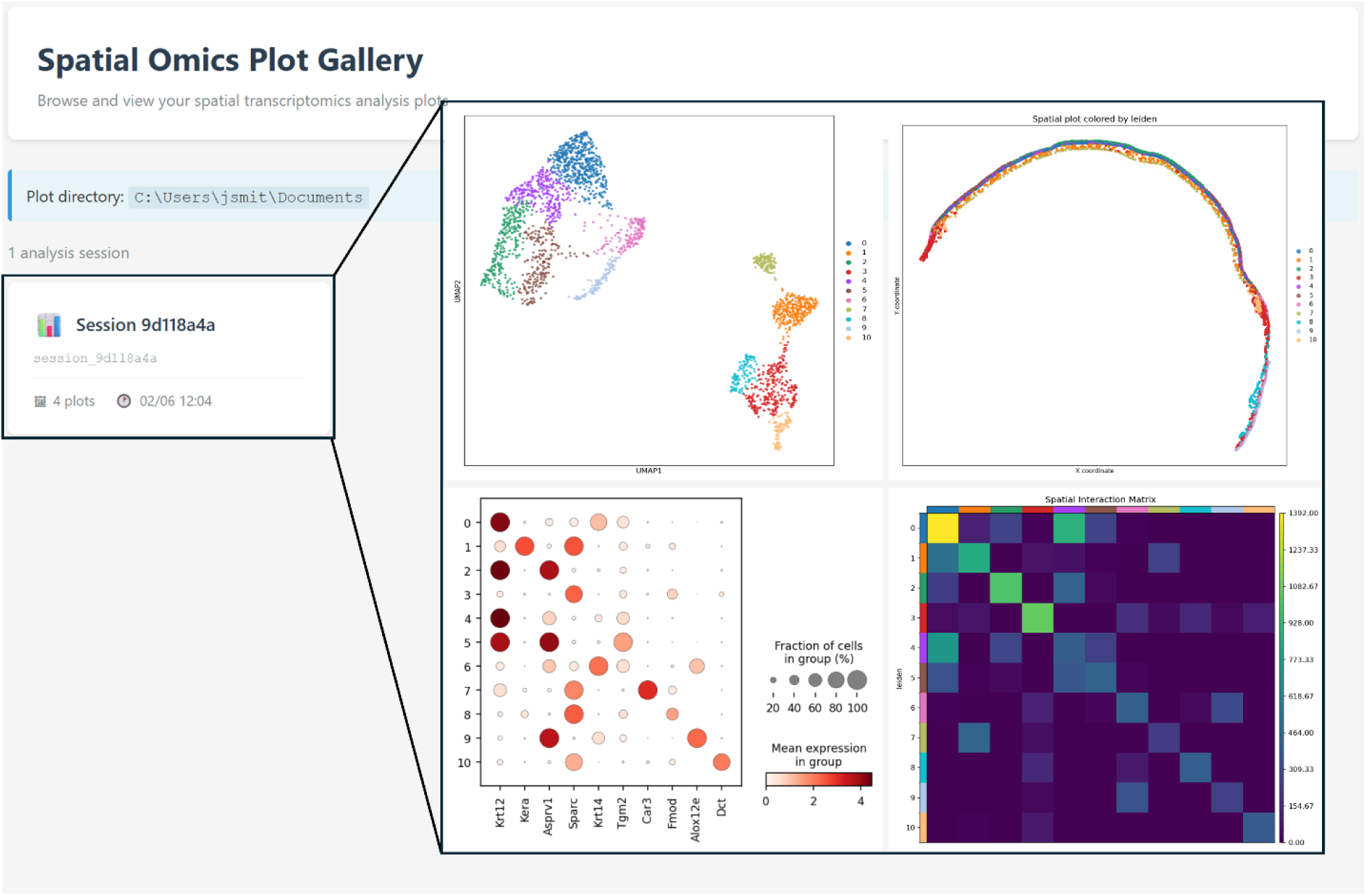
End-user interface for stMCP plot viewer. Web-based plot viewer interface displaying visualizations generated by the MCP server, enabling interactive inspection and organized access to analysis outputs.

#### Deployment and Accessibility

The entire framework is distributed via a public GitHub repository and engineered for fully local, secure execution without containerization, cloud services, or manual environment configuration. All dependencies are resolved during development using optimized Conda-forge binaries and bundled into a portable, self-contained runtime environment. End users install the system through a single executable setup script that automatically extracts the appropriate Python environment configuration, deploys the MCP server modules, and registers the MCP server with Claude Desktop. No manual environment creation, dependency installation, or command-line configuration is required. The MCP server launches locally using standard input/output transport, and all spatial transcriptomics data processing occurs entirely on the user’s machine, ensuring both ease of use for experimental researchers and complete data privacy. Once installed, initiating analysis requires only specification of the directory containing the spatial transcriptomics dataset, enabling immediate exportation without additional setup steps. Simple step-by-step instructions can be found in Supplementary Table 1.

### Biological Significance and Implications for Discovery Analysis

To evaluate the biological utility of the MCP framework, we analyzed a Xenium spatial transcriptomics dataset of mouse corneal tissue (Fig. 4). The corneal dataset comprises predominantly epithelial, stromal, endothelial, immune, and stem cell populations.^33,34^ Within this cellular landscape, certain populations of cell types, such as Schwann cells, are expected to occur at a very low frequency and may be underrepresented or obscured in standard clustering-based analyses due to their rarity.

**Figure 4:**
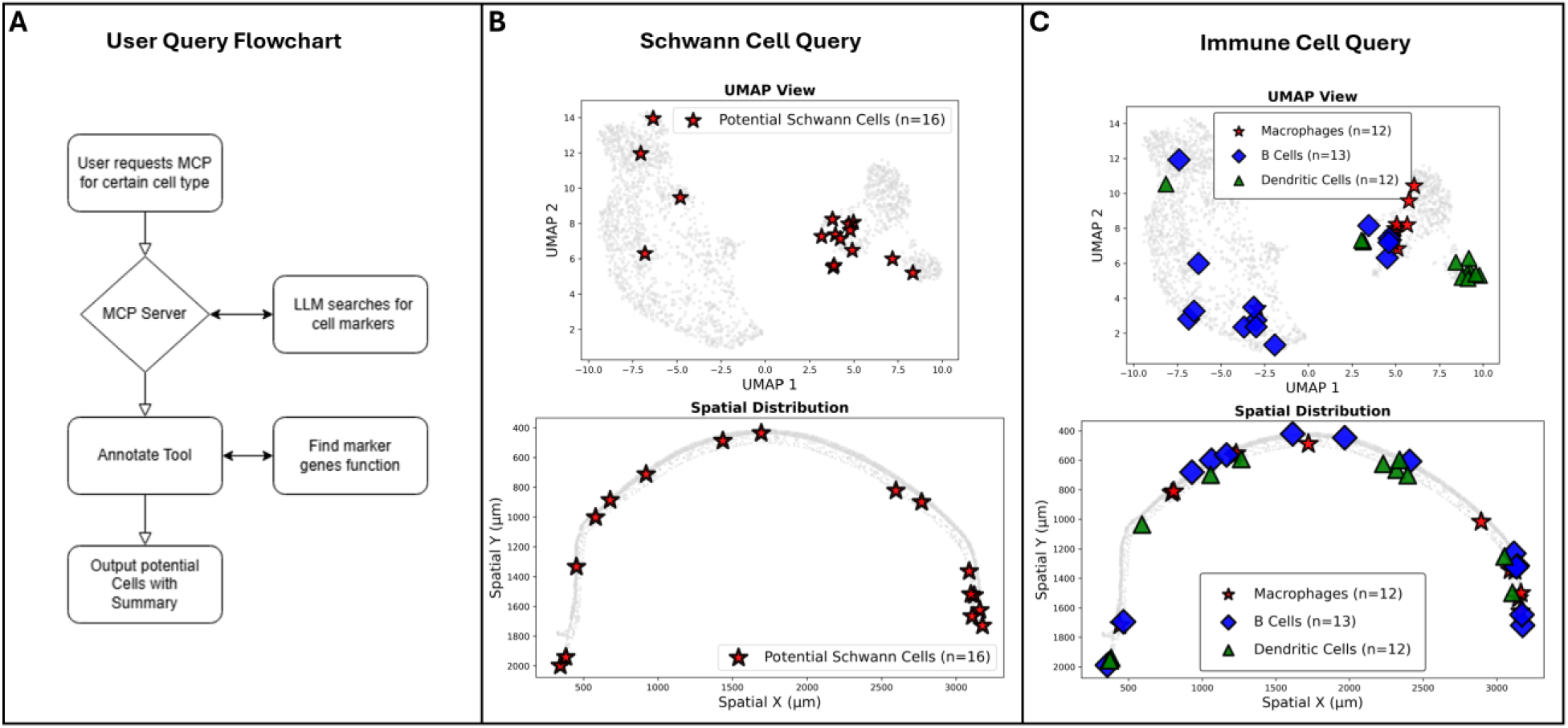
Schwann Cell User Query Flowchart. (A) Flowchart depicts how the MCP server can find hidden cell type populations using both the LLM knowledge base and tool call executions. (B) Example UMAP and spatial plot of potential Schwann cells indicated by the MCP server. (C) Example UMAP and spatial plot of candidate immune cell groups.

Using MCP guided querying, we identified rare cells displaying concordant expression of multiple established Schwann cell-associated marker genes. Structured interrogation of marker co-expression patterns enabled identification of approximately 16 candidate Schwann cells distributed along the corneal tissue arc. These cells were not readily distinguishable as a discrete cluster in unsupervised dimensionality reduction, highlighting the utility of targeted, knowledge-guided exploration layered onto conventional analysis outputs. Experts in corneal immunology confirmed that at least some of the candidate cells were likely Schwann cells and agreed that the selected targets were well justified. While the LLM-identified markers (*Sox10*, *S100b*) are shared by both Schwann cells and melanocytes, the latter are restricted to the corneal periphery; consequently, candidate cells within the central cornea are almost certainly Schwann cells.^35^

In addition to rare cell detection, MCP-guided workflows facilitated refinement of immune cell annotations within the corneal microenvironment. By iteratively querying lineage defining markers and examining their spatial distribution, the framework supported discrimination of immune subsets, including macrophage, B cell, and dendritic populations. Corneal immunology experts agreed that the markers used for macrophages (*Adgre1*, *Csf1r*, *Ptprc*) and dendritic cells (*Zbtb46*) allow for justified labelling by the LLM. The markers used for B-cells (*Cd79b, Ighm*) could support B-cell labeling, but are expressed in low levels and B-cells are not typically found in the central cornea during homeostasis, indicating further investigation may be needed.^36^ This demonstrates that the MCP framework is not limited to rare cell discovery but also supports the detailed delineation of biologically relevant subpopulations within heterogeneous cellular compartments.

Although definitive validation of specific cell identities requires independent experimental confirmation, these examples illustrate how constrained natural language orchestration can integrate prior biological knowledge with reproducible computational workflows. Rather than replacing unsupervised analysis, MCP augments it by enabling iterative, hypothesis-driven interrogation of marker co-expression and spatial organization within a single session.

This capability is broadly applicable across spatial transcriptomics studies, particularly in tissues and disease states where rare, transitional, or spatially restricted cell states may play critical functional roles. By lowering friction between biological hypothesis generation and computational execution, the MCP expands the practical scope of discovery analysis while preserving methodological rigor and traceability.

### Performance and Benchmarking of the MCP Orchestrator

To quantitatively evaluate the MCP orchestrator, we constructed a dataset of 125 natural language prompts reflecting realistic spatial transcriptomics workflows (e.g., data loading, clustering, differential expression, and spatial statistics). (Supplemental Excel Data) Prompts were categorized into single-step, multistep, and ambiguous/invalid requests (Supplementary Table 2). Valid prompts contained sufficient information to satisfy tool schema requirements, whereas invalid ones intentionally lacked mandatory parameters or requested unsupported operations.

Across all evaluated categories, the orchestrator achieved an overall success rate of 95%. For valid requests, successful execution was defined strictly as correctly interpreting user intent, routing to the appropriate tool, assigning valid parameters, generating the expected outputs, and maintaining session state with proper logging. Conversely, ambiguous or incomplete prompts were evaluated for recoverability rather than immediate failure. In 91% of these specific edge cases, the orchestrator successfully preserved session integrity by requesting clarification or proposing valid alternative actions consistent with the available toolset, guiding the user back toward an executable workflow.

Collectively, these results demonstrate that the MCP orchestrator enforces structured tool usage while remaining robust to underspecified input. Importantly, this constrained orchestration model significantly reduces the risk of unintended code execution or uncontrolled analytical branching frequently observed in unconstrained autonomous agent systems.

### Performance and Benchmarking of the MCP Tools

To characterize the computational efficiency of the MCP server, we benchmarked both language model token usage and local execution performance across multiple spatial transcriptomics analysis tasks (Figure 5A). Because the MCP server is explicitly designed to offload all data intensive computation to locally executed tool modules, language model usage and computational runtime represent complementary but distinct components of the overall system performance. The test dataset used consists of about 2,500 cells and 5,100 genes.

**Figure 5:**
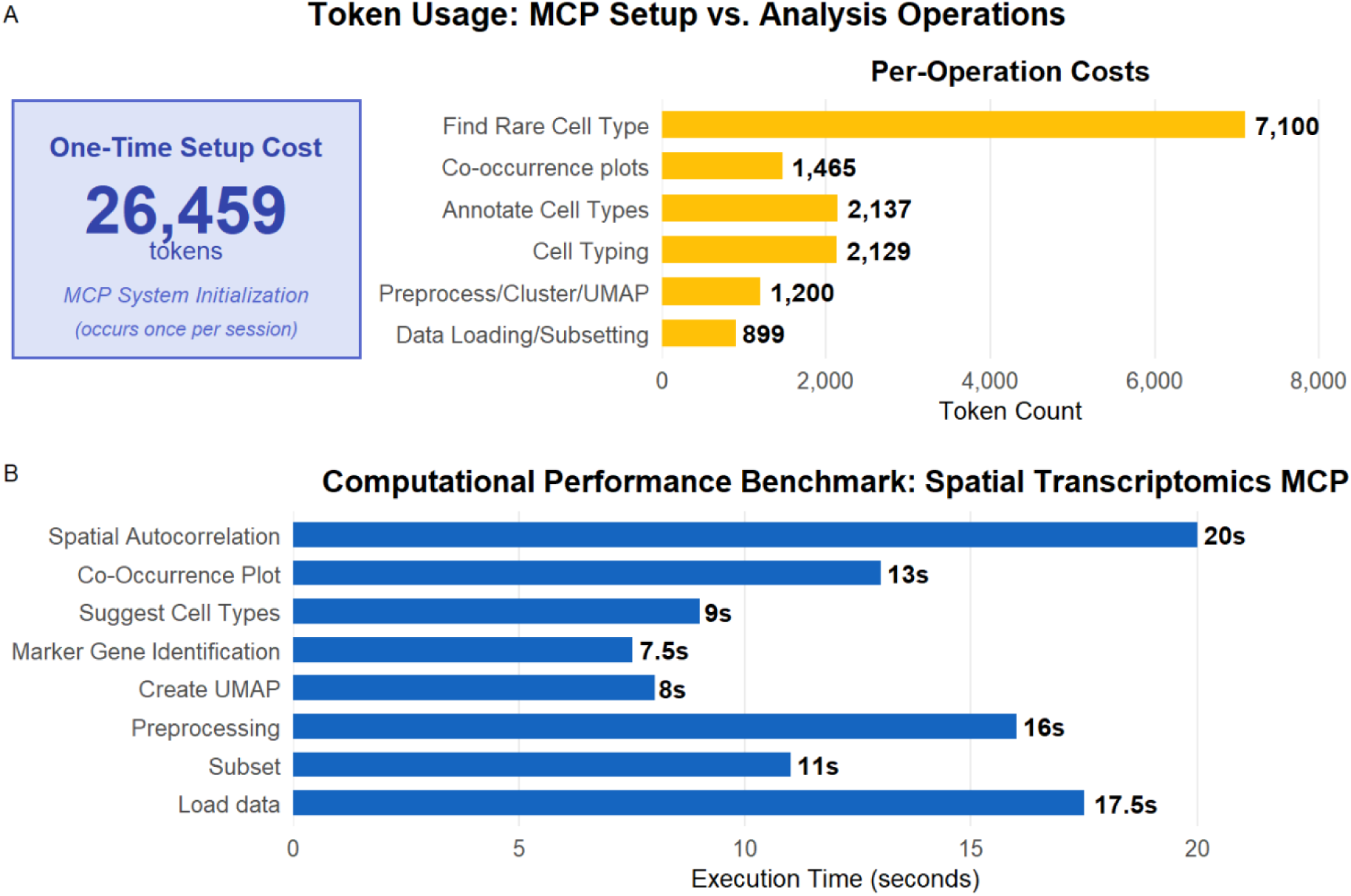
MCP Token and Time Execution Benchmarking. Token consumption and execution time were measured under representative conditions to provide users with practical expectations for MCP performance. This performance may change due to internet speed, hardware specifications, and server connection. (A) Displays token usage of the setup and per-operation costs for specific tool calls. (B) Displays time execution for some common tools calls.

Token usage analysis revealed a one-time initialization cost associated with establishing the MCP session, after which per-operation token consumption remained low. Importantly, token usage is largely independent of dataset size and instead depended primarily on the length of the user’s query and the ensuing dialog between the user and LLM. Analytical computations were executed locally and therefore did not contribute to LLM token costs. Also, Claude now has a one million token context window; therefore, users using all available tokens in a single session are highly unlikely.

In parallel, we measured local execution time for representative analysis steps using a standardized lab workstation configuration (Figure 5B). Execution time scaled with dataset size and task complexity, consistent with conventional spatial transcriptomics workflows, and MCP orchestration introduced minimal additional overhead beyond underlying tool execution. The specifications of the user’s individual workstation and their internet speed will affect execution time. This test was done using a laptop running Windows 11 Pro with an Intel Core i7-14700HX CPU @ 2.10 GHz and 32 GB RAM. Together, these benchmarks provide users with practical expectations for both analysis latency and language model usage when deploying our MCP workflow.

By decoupling language-driven orchestration from numerical computation, the MCP achieves efficient use of LLM resources while preserving interactive, exploratory analysis. This integrated benchmarking demonstrates that MCP spatial transcriptomics analysis is both computationally tractable on local hardware and cost efficient in terms of LLM usage, supporting its suitability for routine exploratory and discovery research.

## Discussion

In this study, we developed and evaluated an MCP framework for reproducible, natural language guided spatial transcriptomics analysis. The MCP enables flexible exploratory analysis while leveraging natural language processing of the LLM. We demonstrated the biological utility of this approach by identifying a rare cell type population within our Xenium dataset and benchmarked the framework for orchestration robustness, language model token efficiency, and local execution performance. Together, these results illustrate how MCP workflows can augment conventional analysis, empowering rather than replacing biological expertise and established computational methods.

### MCP as an Assistive Framework

The interpretation of spatial transcriptomics data fundamentally relies on biological knowledge, experimental context, and careful human validation. Datasets typically comprise expression measurements for thousands of genes across tens to hundreds of thousands of spatially resolved cells, creating a high-dimensional search space where meaningful signals can be difficult to isolate.^37,38^ In manual workflows, researchers are often forced to make early, hypothesis-driven decisions that can inadvertently bias downstream exploration.

By contrast, the MCP functions as an assistive framework that helps structure complex analyses, manage analytical state, and surface candidate signals. It allows users to iteratively explore datasets through natural language queries while preserving full reproducibility, enabling the interrogation of alternative hypotheses without committing to a single rigid workflow. Importantly, the system is not intended to independently generate biological conclusions; rather, it facilitates the efficient navigation of complex datasets to identify patterns that merit deeper expert investigation.

### Advantages and Limitations Relative to Manual Coding and Agent Systems

Compared with manual coding workflows, the MCP lowers technical barriers to spatial transcriptomics analysis by eliminating the need to type code, manage libraries, and learn codebase systems. This approach can reduce development time, support rapid exploratory analysis, and make advanced spatial analyses more accessible to researchers without extensive programming experience. Unlike workflows in which users prompt a general-purpose LLM to generate analysis scripts, MCP executes predefined tool modules rather than dynamically generated code. This design reduces variability across sessions, avoids silent errors introduced through ad hoc code synthesis, and ensures that identical inputs produce reproducible analytical outputs. At the same time, because MCP tools wrap established libraries, users can retain access to validated computational methods rather than opaque model generated computations.

Relative to autonomous AI agent systems, the MCP server adopts a deliberately constrained design. While agent approaches often embed reasoning, planning, and execution within a single mode loop, the MCP separates language understanding from computational execution.^24–26,28^ Analytical operations are performed exclusively by deterministic, locally executed tools, while the LLM serves as an interface for intent specification and orchestration. Also, agentic systems can rely on API external endpoints, that when updated, may cause downstream analysis problems. Meanwhile, MCP tool modules are contained to a local server, allowing updates to increase its capabilities without disrupting future analysis. This separation improves transparency, reduces language model costs, and enables auditing of each analytical step.

Importantly, because all numerical computation occurs on the user’s local machine, MCP workflows avoid the recurring cloud compute costs typically associated with agent-based systems that execute analysis remotely. Language model usage is restricted to intent parsing and output summaries, minimizing token consumption following session initialization.

Nevertheless, MCP workflows remain dependent on the quality of user input and prior biological knowledge. As with any interactive analysis system, poorly specified queries or low-quality data may lead to misleading results. The MCP mitigates but does not eliminate the need for careful experimental interpretation and validation from the user.

### Implications for Biological Discovery in Spatial Transcriptomics

The application of this framework to corneal spatial transcriptomics data illustrates its potential to facilitate discovery, particularly for rare or spatially restricted cell populations. By enabling flexible marker exploration combined with immediate visual feedback, MCP workflows help researchers identify subtle biological signals that might be overlooked in rigid, conventional pipelines.

An additional strength of the MCP architecture is its capacity to incorporate domain-specific "skills."^39–41^ In the broader AI ecosystem, skills and modular knowledge bases, often implemented via retrieval-augmented generation (RAG) or custom plugins, have become standard mechanisms for grounding general-purpose LLMs in specialized domains. By adopting this paradigm, future iterations of our framework could seamlessly integrate curated marker gene lists, lineage hierarchies, or tissue-specific ontologies as discrete modules. Because these knowledge bases can be added without altering the core computational server, the framework is well-positioned to evolve from a computational orchestrator into a biologically grounded analytical assistant.

### Reproducibility as a Design Principal

A central goal of this framework is to make reproducibility an intrinsic, effortless property of spatial analysis. The MCP automatically records user prompts, tool calls, parameter overrides, and generated outputs (Supplementary Fig. 2). The logging captures both the analytical intent and the exact computational execution, enabling workflows to be perfectly reproduced, audited, and shared without manual reconstruction. By preserving context alongside results, the system fosters transparency in exploratory analyses where intermediate decisions heavily influence downstream interpretation.

### Educational and Collaborative Impact

In addition to its analytical capabilities, the MCP has the potential to serve as an educational and collaborative interface for spatial transcriptomics research. By enabling users to express analytical goals in natural language while exposing underlying computational steps, the MCP can help trainees and non-specialists better understand spatial workflows and logic.

Moreover, MCP analysis provides a shared interface through which scientists and bioinformaticians can collaboratively explore datasets, iteratively refine hypotheses, and review analysis decisions. This shared framework may reduce communication barriers and promote more integrated, multidisciplinary approaches to spatial omics research.

### Current Scope and Future Directions

In its current implementation, the MCP supports imaging-based spatial transcriptomics datasets (Xenium, Vizgen MERSCOPE, CosMx Nanostring) and is deployed through Claude Desktop using Claude LLM.^3,29,30^ While this configuration demonstrates the feasibility and utility of the approach, the MCP architecture is intentionally model-agnostic and extensible. As additional LLMs continue to adopt intuitive MCP specifications, this workflow could be extended to alternative LLM backends without changes to underlying tool infrastructure.

The present implementation prioritizes Python-based spatial transcriptomics libraries to maintain a unified runtime environment and deterministic tool execution. Although R-based frameworks such as Seurat are widely used and provide extensive functionality for spatial analysis, integration of cross language tool execution within an MCP framework remains technically feasible but comparatively early in ecosystem maturity.^17^ Managing bidirectional state synchronization, dependency resolution, and reproducibility across heterogeneous runtime environments introduces additional architectural complexity that is currently beyond the scope of this tool. Future work may incorporate R-based analytical modules as technology advances.

Ultimately, the strict separation of orchestration and computation ensures the framework can scale seamlessly, incorporating new multi-omics modalities and analytical methods as the field of spatial biology continues to advance.

## Conclusion

In summary, the spatial transcriptomics MCP framework represents a significant step toward democratizing complex biological data analysis. By bridging the gap between intuitive natural language reasoning and rigorous, deterministic computation, the system empowers biologists to rapidly explore high-dimensional tissue data without compromising reproducibility, data privacy, or incurring prohibitive cloud-computing costs. Beyond spatial biology, the adoption of MCP-based architectures offer a transformative path for the broader scientific community. By providing a standardized, secure interface for connecting AI models to domain-specific analytical engines, MCP can be adapted across diverse fields, from materials science to proteomics, to create a stable, transparent, and expert-aligned research environment. As spatial omics technologies scale in resolution and complexity, cost-effective and transparent frameworks will be essential. By seamlessly marrying computational power with human expertise, these tools will accelerate the next generation of biological discovery.

## Methods

### Dataset and Tissue Preparation

Spatial transcriptomics data were generated from corneal tissue acquired from fifteen 3-month-old mice. The cohort consisted of eight Awat2^-/-^ mice, a model for evaporative dry eye disease, and seven wild-type controls.^42^ Tissue sections were processed for spatial profiling using the Xenium platform (10x Genomics) according to the manufacturer’s instructions, utilizing a 5,106-gene targeted transcript panel.^29^ Following imaging and onboard cell segmentation, expression matrices and spatial coordinates were generated. For biological discovery and framework benchmarking, the dataset was refined to include only corneal tissue regions.

### MCP System Architecture and Implementation

The MCP framework was implemented in Python and executed within a Conda-managed environment. The core analytical engine utilizes Scanpy and Squidpy, supported by the standard scientific Python stack (NumPy, Pandas, and Matplotlib).^18,19,32,43,44^ The MCP server functions as a local backend process interfaced through the Claude Desktop client (Anthropic).

Communication between the host and server is mediated via structured Model Context Protocol (MCP) tool invocations. Crucially, the large language models (Claude Sonnet and Opus) are restricted to interpreting user intent and generating natural language summaries; all numerical computation, data transformation, and visualization are executed locally. All analyses were performed on a workstation running Windows 11 Pro (Intel Core i7-14700HX CPU @2.10 GHz, 32 GB RAM) without GPU acceleration.

### Orchestrator Benchmark Evaluation

To quantitatively assess the orchestrator, we benchmarked its performance against a curated set of natural language prompts simulating common spatial transcriptomics workflows. These prompts spanned the full library of available tools and included complex, multi-operational sequences. We categorized inputs into three tiers: single-step, multistep, and ambiguous or invalid. Valid prompts provided all parameters necessary to satisfy underlying tool schemas, while invalid or ambiguous inputs intentionally omitted required data or requested operations beyond the framework’s scope.

Task completion was rigorously defined by five criteria: (1) accurate intent-to-tool routing, (2) valid parameter mapping, (3) successful execution of the underlying Python calls, (4) generation of the expected analytical outputs, and (5) persistent maintenance of the session state. For ambiguous inputs, success was measured by the orchestrator’s recoverability, specifically its ability to request missing parameters or propose valid alternatives. For invalid requests, performance was evaluated based on the system’s ability to correctly identify and reject unsupported tasks while preserving session integrity. All prompts are stored within the Supplementary Excel Dataset.

### Token Usage Quantification and Runtime Benchmarking

To evaluate the operational efficiency of the framework, we recorded language model token consumption via the Claude interface metrics. Initialization overhead was defined as the total tokens required to establish the MCP session context. Per-operation cost was calculated as the incremental token consumption for individual analytical interactions following initialization. These measurements were made across representative spatial transcriptomics tasks to determine average usage patterns.

Local computational performance was assessed by measuring the execution latency of core MCP tool calls, including preprocessing, clustering, visualization, and spatial statistics. We utilized wall-clock timing on the local workstation to isolate execution time, explicitly from the user entering their request to when the LLM responds with a successful completion message.

### Rare Cell and Immune Subset Identification Criteria

Potential Schwann cell populations were identified by querying the MCP framework for established lineage markers. Cells were nominated as candidate Schwann cells based on the concordant expression of these markers exceeding normalized thresholds established during the preprocessing workflow. Similarly, potential immune cell subsets, including macrophages, B cells, and dendritic populations, were identified through iterative querying of lineage-specific genes. To support these provisional annotations, we leveraged the MCP session context to evaluate gene co-expression patterns in conjunction with spatial localization. Candidate cell populations were subsequently reviewed by domain experts familiar with corneal biology, who independently assessed marker concordance and spatial distribution patterns and agreed that these cells represent plausible targets for further investigation. These identities remain computationally inferred annotations provided to illustrate the utility of the MCP framework; definitive validation would require orthogonal experimental confirmation.

## Code and Data

Code and directions for the setup of the MCP server will be available on GitHub at the time of publication. (https://github.com/jsmithcm10/stMCP)

## Supporting information

Supplemental Prompt Sheet

## Acknowledgements

We thank Dr. Yehe Liu for early conceptual discussions that contributed to the initiation of this work. Spatial transcriptomics data was generated with support from the CWRU Imaging Core. Generative artificial intelligence tools were used to assist with code debugging and language editing during manuscript preparation. All analytical decisions, data interpretation, and final manuscript content were reviewed and verified by the authors.

## Funding

National Institutes of Health (U01-EY034693, UO1-EY034687, R01-HL126747, T32-EB007509, P30-EY011373)

## Notes

### Competing Interest Statement

The authors have declared no competing interest.

