## Supplemental Figures and Tables for "stMCP: Spatial Transcriptomics with a Model Context Protocol Server"

**Supplementary**

Supplemental Figure 1

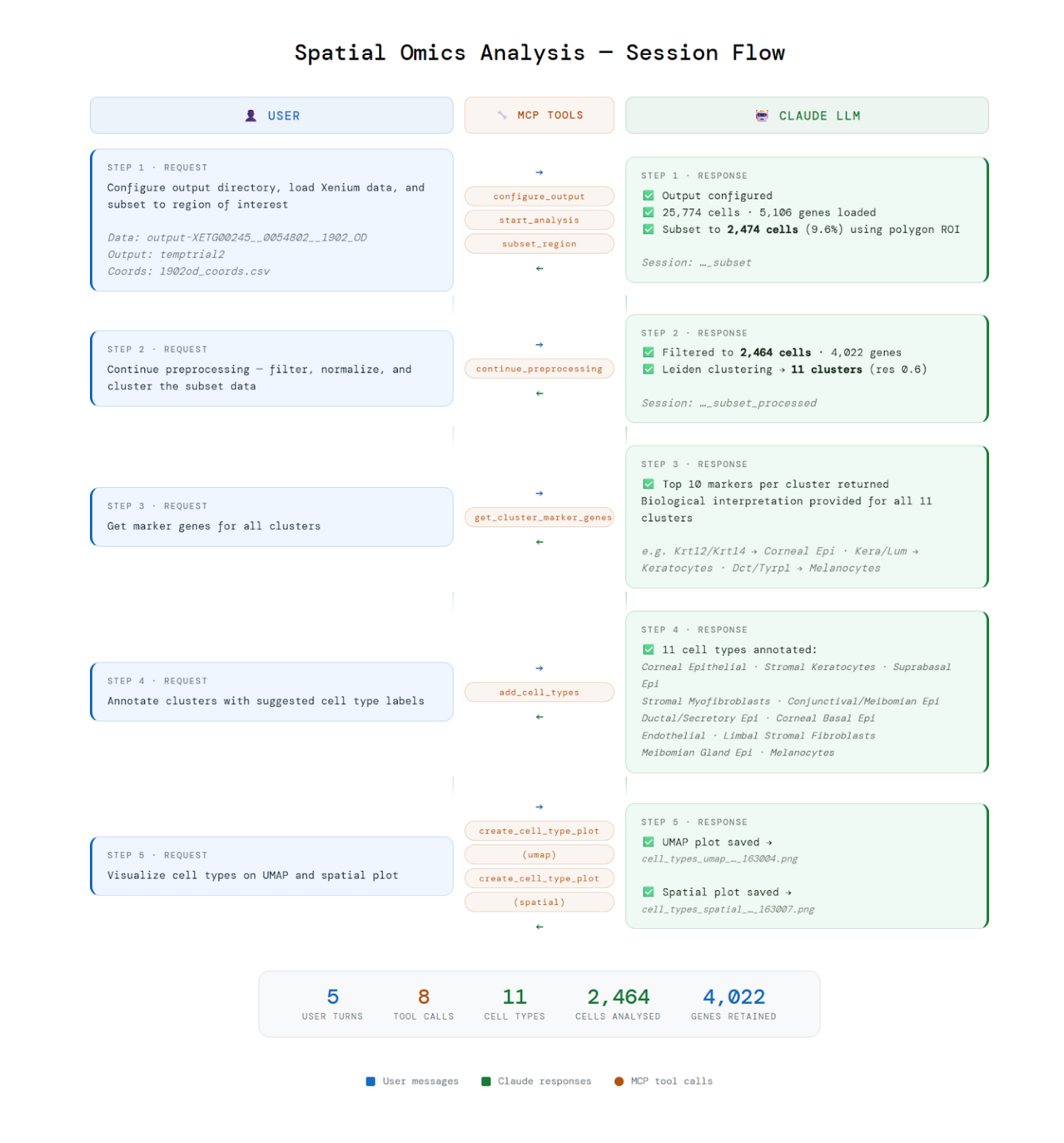

**Supp Fig 1:** **Spatial Transcriptomics Analysis Session Example**: Example conversation between user (blue) and LLM (green) for a selection of tool calls (orange). The user requests are routed, and tools are selected by the orchestrator. These tools are executed and the LLM provides a response with a summary of the request along with other information.

Supplemental Figure 2:

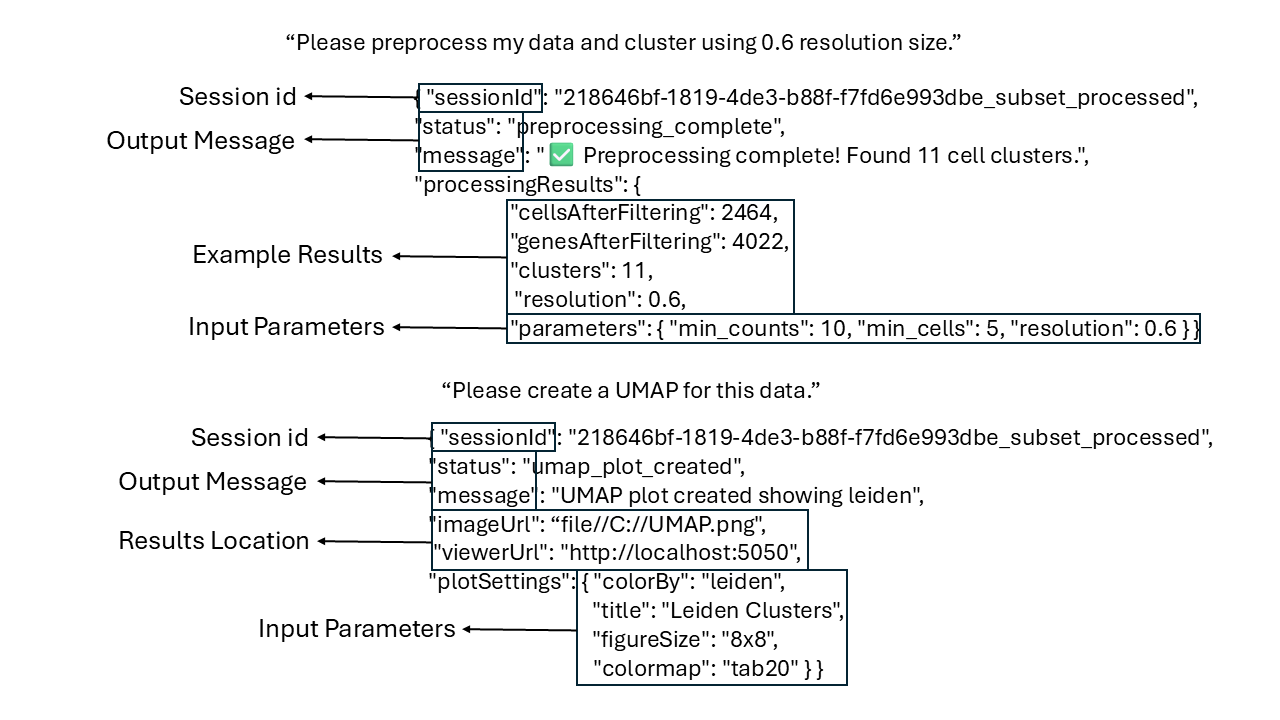

**Supp Fig 2:** **Example Tool Call Logging.** Example of how the MCP logs individual tool calls with two prompts (preprocess/cluster, UMAP generation). This shows the session ids, output message for the LLM, example results that the LLM may produce, locations of outputs, and the input parameters that were needed by the MCP server to execute the code.

Supplementary Figure 3:

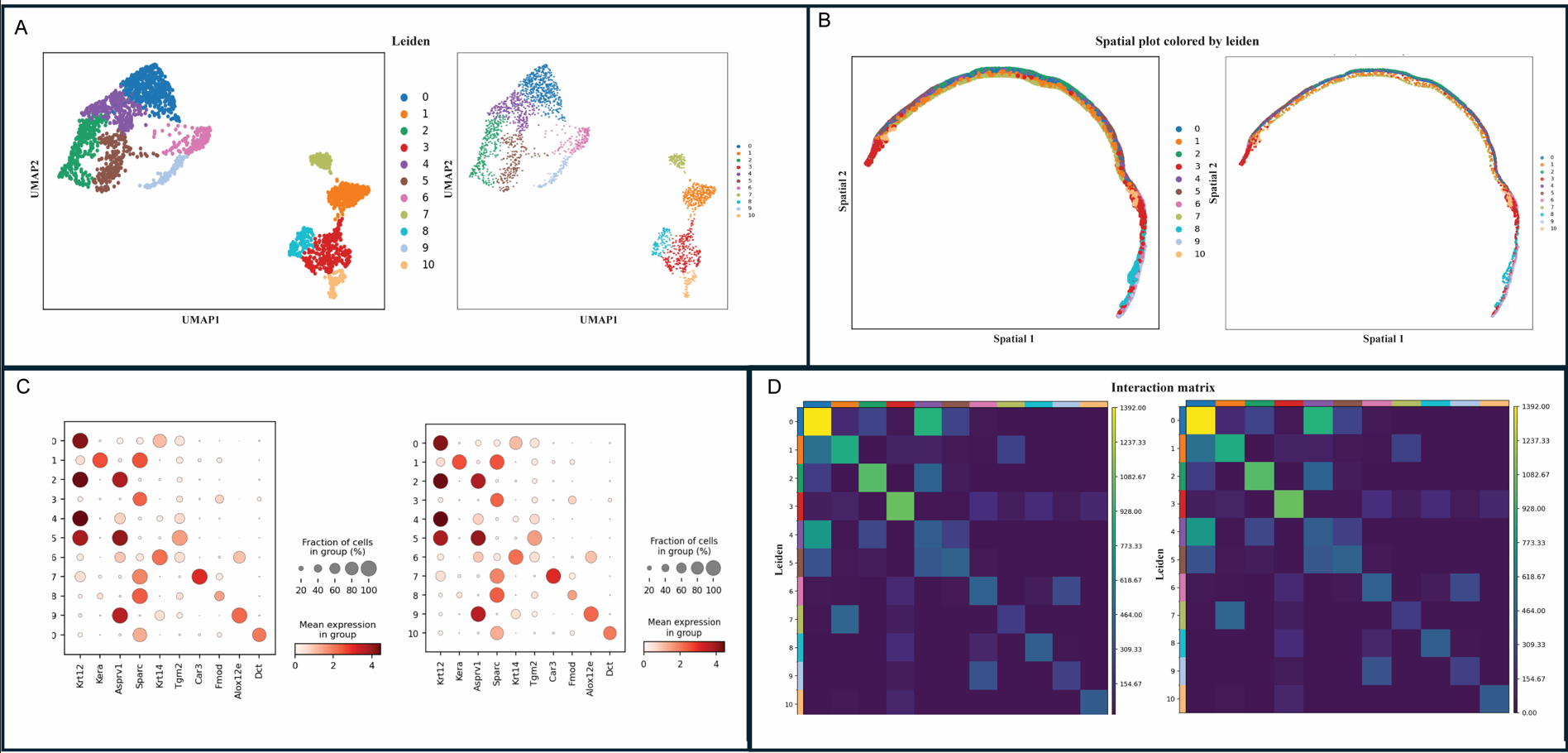

**Supp Fig 3:** **Example of Duplicate Results from Manual Coding and MCP**. Depicts the same results from manual coding (left) and MCP server generated outputs (right) for (A) UMAPs, (B) Spatial plots, (C) Dot plots, and (D) Spatial interaction matrices.

Supplemental Figure 4:

Orchestrator Code:

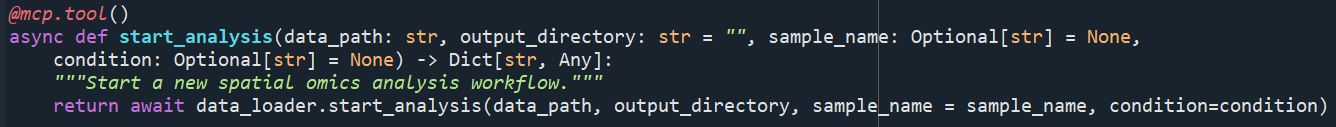

Tool Module:

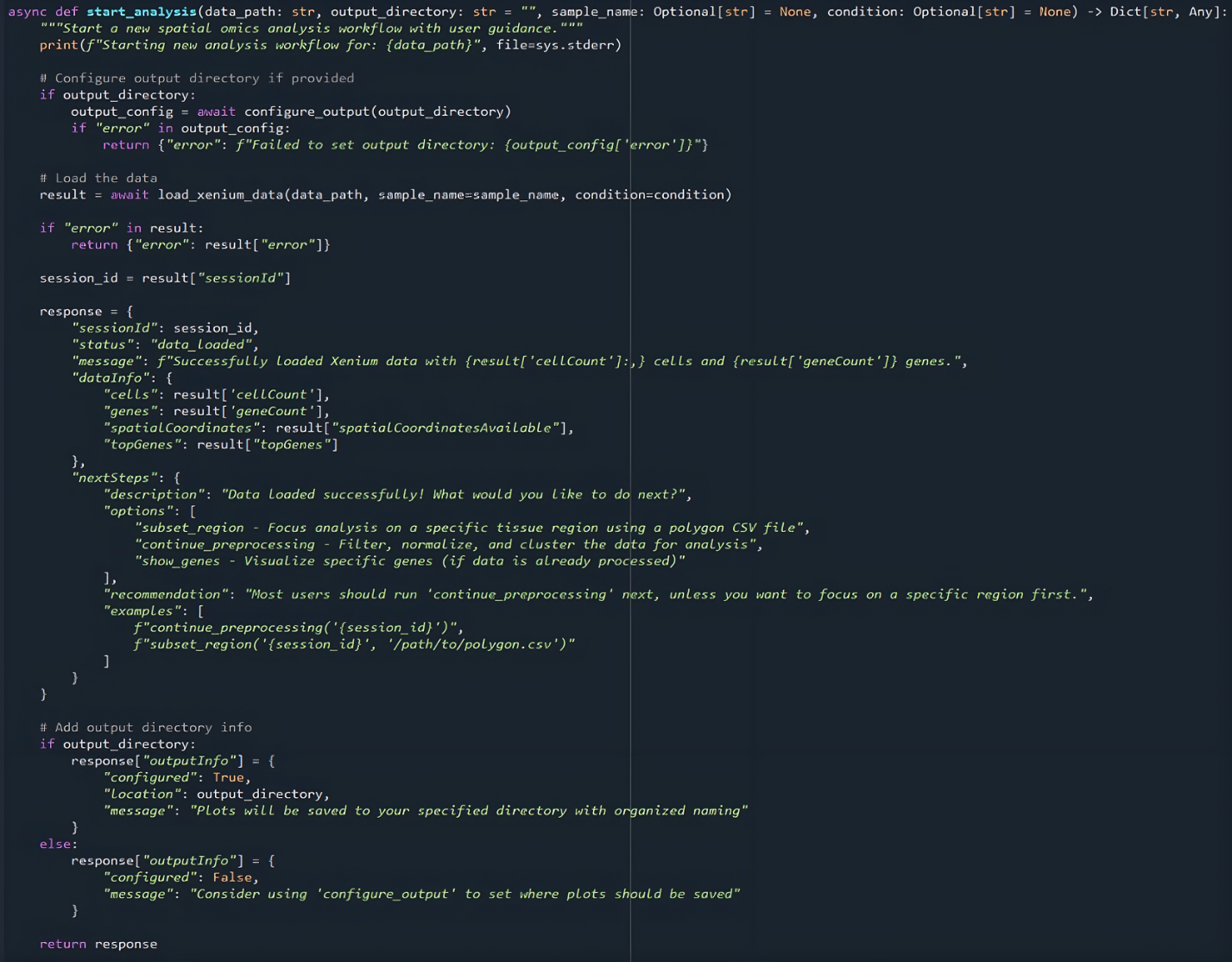

Actual Function:

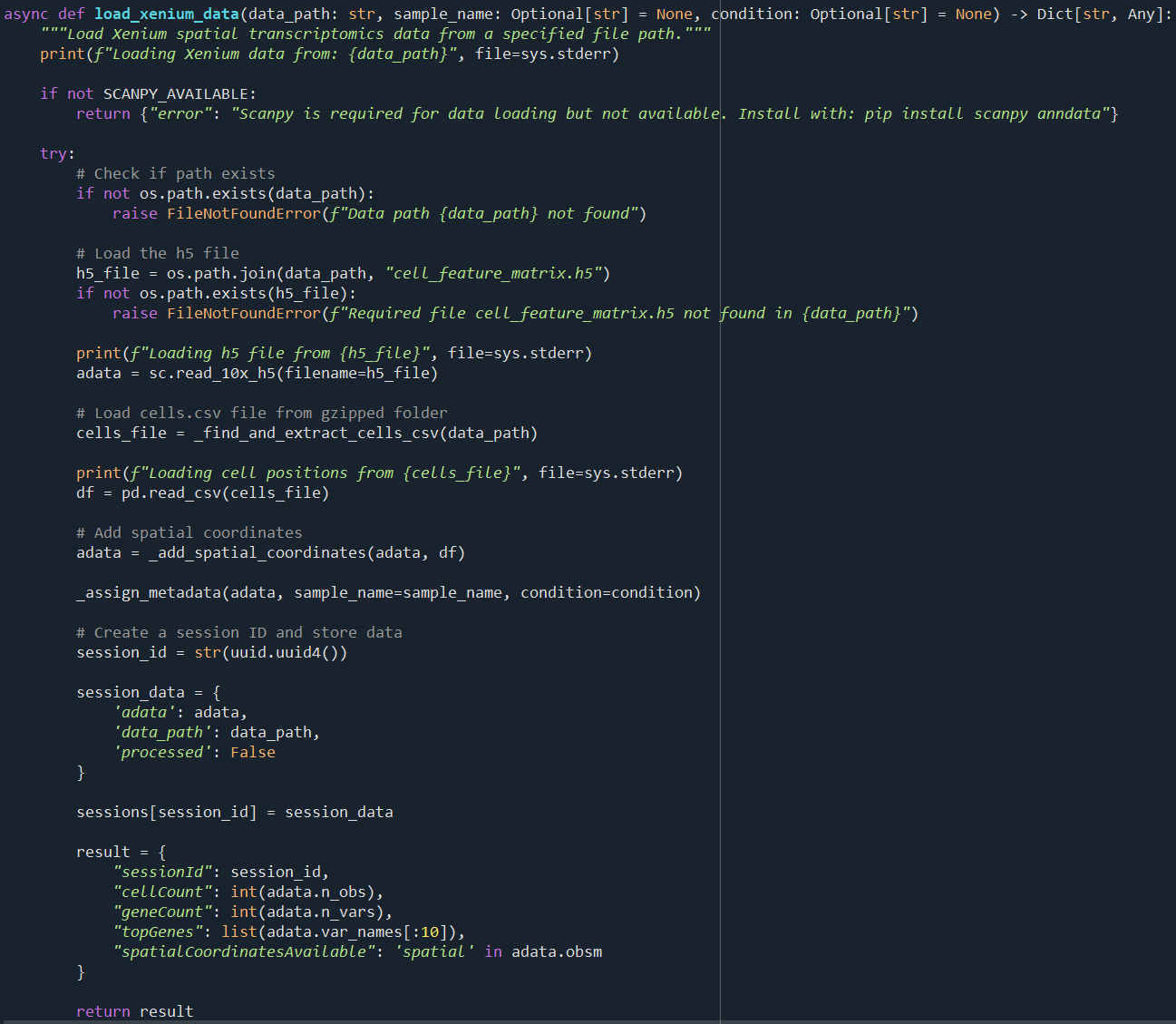

**Supp Fig 4:** **Example Function Wrapping Code.** Depicts an example code snippet for the data loading tool module, specifically loading Xenium data. The top image shows orchestrator code, the middle image the data loading (start analysis) tool module and the final window shows the actual function.

Supplementary Table 1:

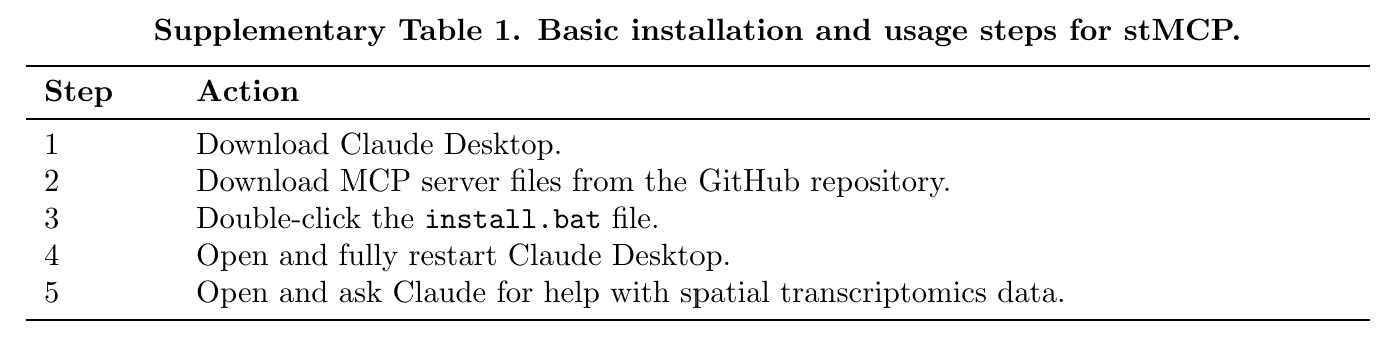

**Supp. Table 1: Installation Instructions.** Simple step-by-step instructions on how to install and start using stMCP.

Supplementary Table 2:

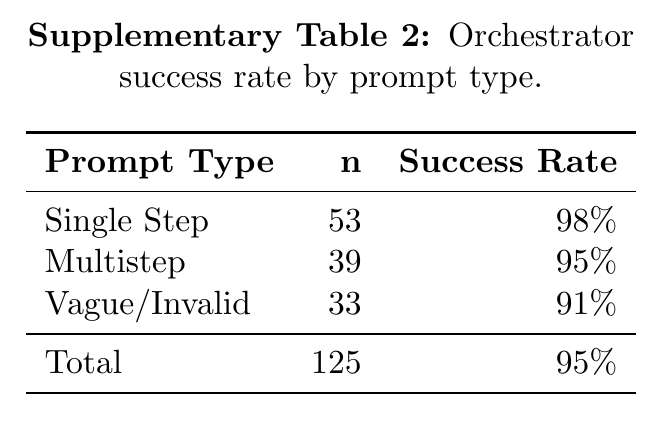

**Supp. Table 2: Orchestrator success rate by prompt type.** Table displays the three prompt types, number of entries per type and success rate as a percentage.
